# Modeling the Growth of Emotional and General Vocabulary Using Nonlinear Trajectories

**DOI:** 10.1101/2025.06.24.661340

**Authors:** J. Formoso, A. D. Calero, D.I. Burin, P. Vazquez-Borsetti

## Abstract

Understanding how emotional and general vocabulary develop across the lifespan offers key insights into cognitive and socioemotional processes. While emotional vocabulary is foundational for emotion regulation and social interaction, general vocabulary underpins broader cognitive functions such as reasoning and reading comprehension. In this study, we modeled the growth trajectories of emotional and general vocabulary using Gompertz functions in a large cross-sectional sample (N = 820; age range = 12– 84 years). To control for item difficulty, we selected a subset of vocabulary items with similar and low difficulty in adulthood. Both vocabulary types showed non-linear growth patterns, with emotional vocabulary exhibiting earlier and faster development, reaching near-asymptotic levels by early adulthood. General vocabulary showed a more gradual increase and later inflection point. A joint nonlinear mixed-effects model confirmed significant differences in developmental timing, with emotional vocabulary peaking approximately four years earlier than general vocabulary. These findings support theoretical models emphasizing the early emergence and adaptive relevance of emotional concepts and highlight adolescence as a sensitive period for emotional vocabulary acquisition. The Gompertz model proved effective in capturing asymmetric vocabulary growth, providing interpretable parameters aligned with developmental theory. Implications for education and emotional development interventions are discussed.

## Introduction

Understanding how individuals acquire and refine emotional vocabulary is essential for studying emotion processing and regulation across the lifespan. The Theory of Constructed Emotion (Barrett, 2017a), formerly known as the Conceptual Act Theory, offers a compelling framework for understanding how emotional experiences are shaped through conceptual knowledge and language. According to this theory, emotions are not fixed biological states but rather flexible conceptual categories formed through accumulated experience, in which language plays a central role (Barrett, 2017b; Hoemann, Devlin, et al., 2020; Hoemann, Wu, et al., 2020).

Language enables individuals to identify regularities in the environment, build shared conceptual categories, and organize emotional experiences (Lindquist, 2017). Emotional vocabulary is therefore not merely a linguistic repertoire but a cognitive tool that structures emotional perception and regulation (Shablack & Lindquist, 2019). Within this framework, the construct of emotional granularity—the ability to distinguish and label emotions with precision—has received growing attention with higher emotional differentiation has been linked to adaptive outcomes such as improved emotion regulation (Cole et al., 2010; Holodynski et al., 2013), reduced aggressive responses (Pond Jr. et al., 2012), and better psychosocial functioning (Ciarrochi et al., 2008).

Research on development consistently indicates that emotional vocabulary grows with age, although the trajectory of this growth differs across studies. Some findings show that between ages two and five, children begin to label basic emotions, and that their repertoire of emotion-related words approximately doubles every two years thereafter, stabilizing around age 12 (Baron-Cohen et al., 2010; Grosse et al., 2021). Other studies, however, propose that the expansion of emotion-specific vocabulary continues beyond childhood, with older age associated with greater differentiation and abstraction in emotional language (Bazhydai et al., 2019; Nook et al., 2018, 2020; Shablack & Lindquist, 2019).

Grosse et al. (2021) showed that children’s emotion vocabulary becomes progressively more specific and converges with adult usage throughout development. This acquisition process is shaped by semantic features such as valence and word specificity. Complementarily, Ornaghi and Grazzani (2013) showed that both the comprehension and use of emotional-state language are significantly associated with children’s emotion understanding. Furthermore, a greater capacity to distinguish between emotional states has been associated with better emotion regulation, improved psychological adjustment, and reduced reliance on maladaptive coping strategies (Kashdan, Barrett, & McKnight, 2015). Recent research suggests that emotional vocabulary is shaped by both individual experiences and educational context. For instance, psychology students show greater emotional vocabulary than students from other fields, and this increases with years of university training (Calero et al., 2023).

In parallel, general vocabulary has long been recognized as a foundation for diverse cognitive functions. It contributes to reading comprehension (Ouellette, 2006; Perfetti & Hart, 2008), problem-solving (Cirino et al., 2016), and logical reasoning (Cain & Oakhill, 2014; Kintsch, 1998). Both vocabulary breadth (number of known words) and depth (quality of lexical knowledge) affect how efficiently individuals process and integrate information (Dicataldo et al., s. f.; Oakhill et al., 2015).

Despite this broad evidence base, few studies have compared the development of general and emotional vocabulary simultaneously across the lifespan. The current study seeks to fill this gap, drawing on the theoretical lens of constructed emotion to examine the role of language in shaping emotional knowledge by describing and comparing the developmental trajectories of general and emotional vocabulary from early adolescence through late adulthood.

Modeling vocabulary development across the lifespan requires approaches that can capture non-linear, age-dependent growth patterns. While linear models offer a simplified view, they often fail to reflect the asymmetric nature of cognitive growth, particularly in domains like language (Moore & Bosch, 2009). In contrast, the Gompertz function—a sigmoidal, asymmetric growth model—has been widely used to describe biological and cognitive developmental trajectories due to its flexibility in capturing both early acceleration and late-life plateaus (Buchanan & Cygnarowicz, 1990; Tjørve & Tjørve, 2017; Vázquez-Borsetti, 2024; Wang & Guo, 2024).

Additionally, the Gompertz model offers interpretability aligned with developmental theory: the lower asymptote corresponds to initial capacity (e.g., early vocabulary knowledge), the rate parameter reflects the pace of acquisition, and the upper asymptote denotes saturation or adult-like competence. Given its mathematical flexibility and theoretical plausibility, the Gompertz function is particularly well-suited to model the differential growth of general and emotional vocabulary across the lifespan.

In addition, authentic developmental growth trajectories should not be confounded with a methodological artifact, e.g., the relative difficulty of vocabulary tests. In this sense, for example, if even for adults items in the emotional vocabulary test are overall easier than items in a general vocabulary test, it would not be warranted to conclude that age differences in those items reflect developmental differences, but could also be reflecting how the tests were constructed, or other factors affecting item characteristics. To address this issue, we carried out the analyses on a subset of items (8 from each scale) exhibiting similar and low difficulty in adulthood. Thus, we will analyze whether there are different developmental trajectories for emotional and general vocabulary items that are well known in adulthood.

## Method

### Participants

Participants included N = 820 individuals ranging in age from 12 to 84 years (mean age = 23.32, sd = 15.77), 64% female, from the Metropolitan Area of Buenos Aires. All individuals participated voluntarily and provided informed consent authorizing the anonymized use of their data. For participants under the age of 18, parental or legal guardian consent was obtained prior to the assessment. Data from individuals with reported sensory, neurological, or learning disorders were excluded from the final analyses.

### Procedure

Adolescents were recruited from private secondary schools. After obtaining authorization from the school authorities and parental consent, tasks were administered in person during regular school hours, in 40-minute sessions. Adult participants were recruited through social media platforms, while older adults (>60) were recruited from a community center that offers cognitive wellness workshops. For the older adult group, the Functional Activities Questionnaire (Novacheck et al., 2000) was administered to screen for cognitive impairment, and any participants showing signs of decline were excluded. Both adults and older adults completed the assessments online after providing informed consent for participation.

### Materials

Emotional Vocabulary: this variable was assessed using the Vocabulary subtest of the Emotion Knowledge instrument developed by (Delgado et al., 2017, 2018). This subtest consists of 40 emotion-related words. For each item, participants are asked to select the option (from five basic emotion labels) that most closely matches the meaning of the target word. The task yields a score representing the participant’s emotional vocabulary breadth. The instrument has demonstrated adequate psychometric properties, including evidence of validity and reliability.

General Vocabulary: this variable was measured using the BAIRES-A (Kohan, 2004), a standardized test comprising 34 multiple-choice items across two subscales: Definitions and Synonyms. In the Definitions subscale, participants select the most appropriate definition for a given word from four alternatives. In the Synonyms subscale, they are asked to choose the word most similar in meaning to the target word. We selected 9 items from each subscale. Scores reflect the number of correct responses, with a maximum of 18 points. The BAIRES-A provides robust indicators of construct validity and internal consistency, with a Cronbach’s Alpha of .74 for this sample.

Demographic Questionnaire: participants also completed a brief ad hoc questionnaire to collect demographic information, including gender (female, male, or other), age, and highest education level acquired.

### Data Analysis

We initially selected adult participants aged 30 to 35 years and identified items from each vocabulary scale that showed identical and relatively low difficulty (i.e., accuracy above 75%) within this age group. From each scale—emotional vocabulary (EV) and general vocabulary (GV)—8 items were selected, and all subsequent analyses were conducted using this subset.

The development of emotional vocabulary (EV) and general vocabulary (GV) as a function of age was modeled using Gompertz functions, which are suitable for describing bounded growth trajectories.

The Gompertz function is a three-parameter sigmoid curve defined as:

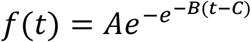

where *A* represents the upper asymptote (expected maximum value), *B* is the growth rate, and *C* is the inflection point corresponding to the age at which the rate of vocabulary growth is fastest. This flexible formulation allows the modeling of accelerated vocabulary acquisition in early stages followed by a deceleration as a theoretical maximum is approached.

Initially, separate nonlinear least squares (NLS) Gompertz models were fitted for EV and GV, estimating the asymptote (A), a parameter related to the growth rate (B), and the inflection point (C) for each vocabulary type. Subsequently, EV and GV scores were normalized by dividing each observation by the corresponding asymptote parameter, allowing for direct comparison on a common scale. A merged dataset was created that included a dummy variable indicating vocabulary type (emotional vs. general).

To assess differences in growth trajectories between EV and GV, an extended Gompertz model was fitted that incorporated interaction terms between vocabulary category and the growth rate and inflection point parameters. This model was first estimated using nonlinear least squares (NLS) and then refined using a nonlinear mixed-effects model (NLME), allowing random variation in the inflection point parameter (C) across individuals to account for interindividual variability.

Additionally, a linear mixed-effects model (LME) was fitted to the normalized vocabulary scores, treating age as a categorical variable and modeling fixed effects of vocabulary type, age, and their interaction, with random intercepts by participant. Post hoc comparisons of EV and GV at each age group were conducted using estimated marginal means (EMMs) with Holm correction for multiple comparisons.

Finally, fitted trajectories, normalized differences between EV and GV, and second derivatives of the growth curves were plotted to visually inspect growth patterns and identify points of maximum acceleration. All analyses were conducted in R version 4.4.1 (R Core Team, 2024), using the packages nlme (Pinheiro & R Core Team, 2023), and emmeans (Lenth, 2023).

## Results

Descriptive statistics for general vocabulary, emotional vocabulary, and age in years for the overall sample are shown in Table 1.

**Table 1.** Descriptive Statistics for General Vocabulary, Emotional Vocabulary, and Age.

| Variable | M | SD | Min | Max |
| --- | --- | --- | --- | --- |
| General vocabulary | 5.19 | 1.99 | 0 | 8 |
| Emotional vocabulary | 5.75 | 1.48 | 1 | 8 |
| Age in years | 23.31 | 15.77 | 12 | 84 |

Note. M = mean; SD = standard deviation; Min = minimum value observed; Max = maximum value observed. General vocabulary and emotional vocabulary scores are presented in raw units.

Separate Gompertz models were initially fitted to GV and EV growth data. Table 2 shows the estimated values for each parameter.

**Table 2.**
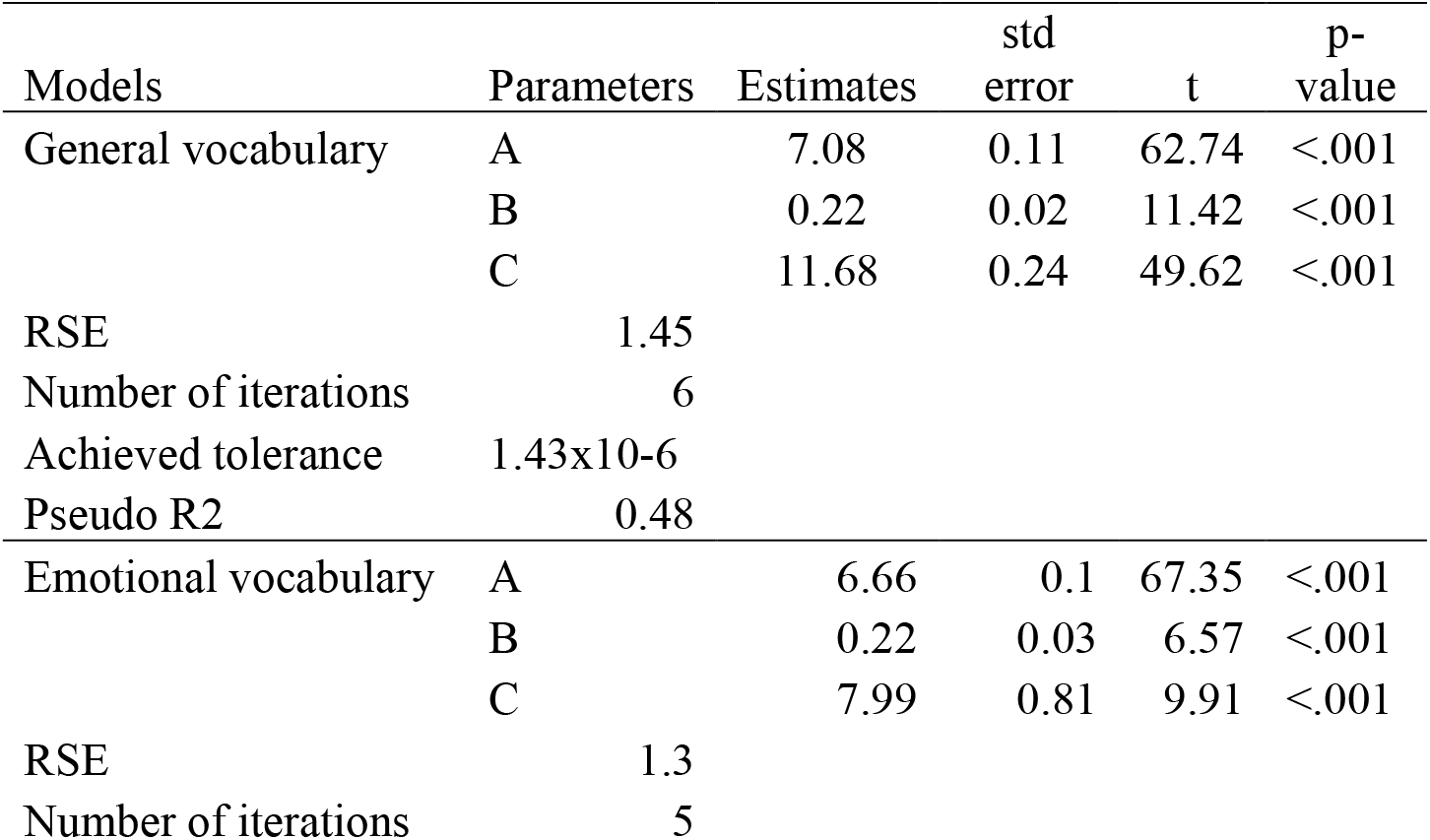
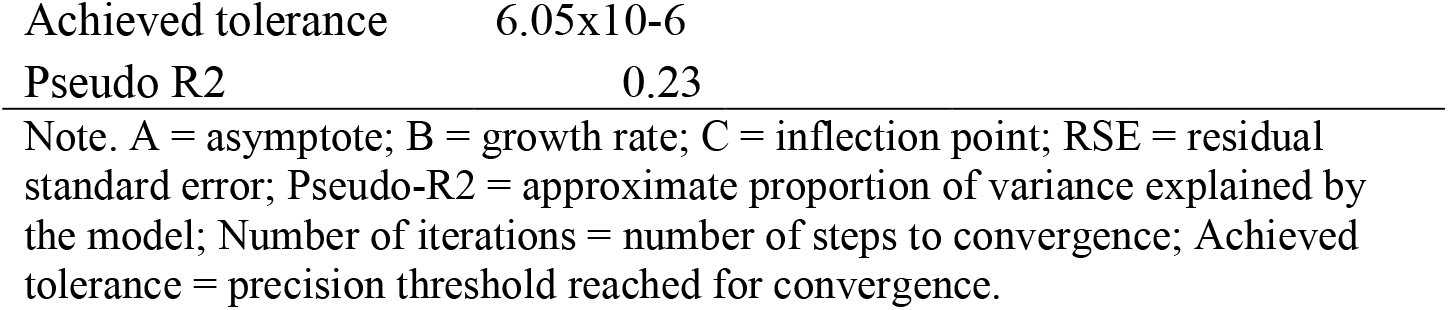
Parameter Estimates and Model Fit Statistics for General and Emotional Vocabulary Growth Models.

| Models | Parameters | Estimates | std error | t | p-value |
| --- | --- | --- | --- | --- | --- |
| General vocabulary | A | 7.08 | 0.11 | 62.74 | <.001 |
|  | B | 0.22 | 0.02 | 11.42 | <.001 |
|  | C | 11.68 | 0.24 | 49.62 | <.001 |
| RSE | 1.45 |  |  |  |  |
| Number of iterations | 6 |  |  |  |  |
| Achieved tolerance | 1.43x10-6 |  |  |  |  |
| Pseudo R2 | 0.48 |  |  |  |  |
| Emotional vocabulary | A | 6.66 | 0.1 | 67.35 | <.001 |
|  | B | 0.22 | 0.03 | 6.57 | <.001 |
|  | C | 7.99 | 0.81 | 9.91 | <.001 |
| RSE | 1.3 |  |  |  |  |
| Number of iterations | 5 |  |  |  |  |

|  |  |
| --- | --- |
| Achieved tolerance | 6.05x10-6 |
| Pseudo R2 | 0.23 |

Note. A = asymptote; B = growth rate; C = inflection point; RSE = residual standard error; Pseudo-R2 = approximate proportion of variance explained by the model; Number of iterations = number of steps to convergence; Achieved tolerance = precision threshold reached for convergence.

The models for general and emotional vocabulary showed statistical significance for all parameters, successful convergence within a small number of iterations, and achieved strict convergence tolerances, indicating stable parameter estimation. The residual standard errors for both models were low, suggesting that the fitted curves closely approximated the observed data. Pseudo-R2 values indicated moderate proportions of explained variance for both vocabulary types, consistent with expected variability in developmental data.

Residual analyses for the general and emotional vocabulary models supported the adequacy of the Gompertz fits. Scatterplots of predicted values against residuals revealed constant variance across the range of predictions, and histograms of residuals indicated approximate normality. These patterns suggest that model assumptions of homoscedasticity and normality were reasonably met.

Visual inspection of the scatterplots (Figure 1) revealed that the observed vocabulary scores closely followed the trajectories predicted by the fitted Gompertz functions for both general and emotional vocabulary.

**Figure 1.**
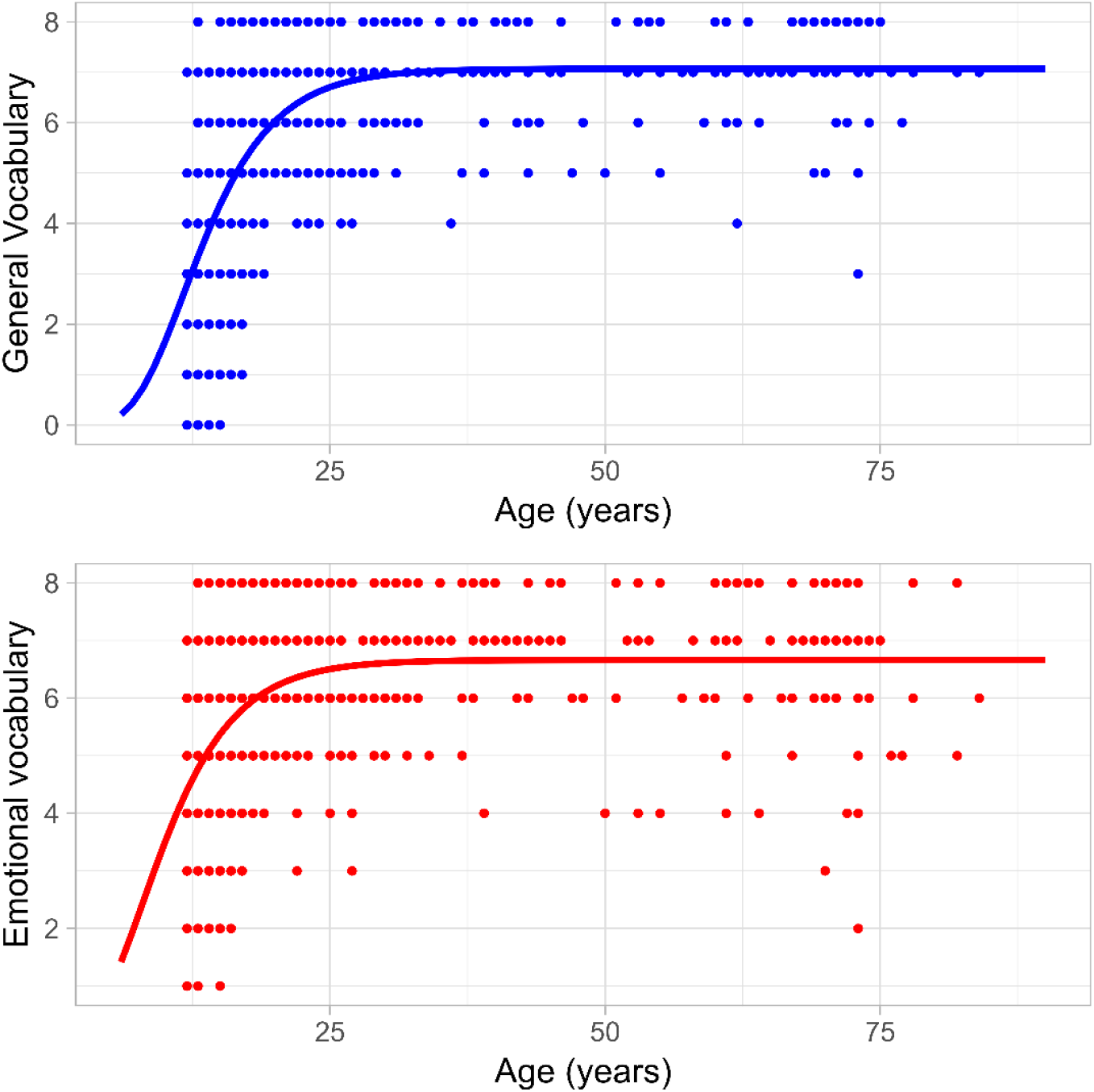
Observed and Fitted Growth Curve for General and Emotional Vocabulary Scores. The scatterplots show observed general (blue dots) and emotional (red dots) vocabulary scores as a function of age, overlaid with the fitted Gompertz curve.

The distribution of points aligned well with the fitted curves across the full range of ages, supporting the adequacy of the models in capturing the underlying growth patterns.

To better understand the dynamics of vocabulary acquisition over time, the second derivatives of the fitted Gompertz curves were computed and plotted (Figure 2). The second derivative reflects the acceleration or deceleration in growth rates, providing insight into the periods of most rapid developmental change. For emotional vocabulary, the peak in acceleration was reached earlier in development (12 years), whereas for general vocabulary, it occurred at a comparatively later stage (16 years). These findings suggest that emotional vocabulary acquisition undergoes an earlier acceleration compared to the development of general vocabulary.

**Figure 2.**
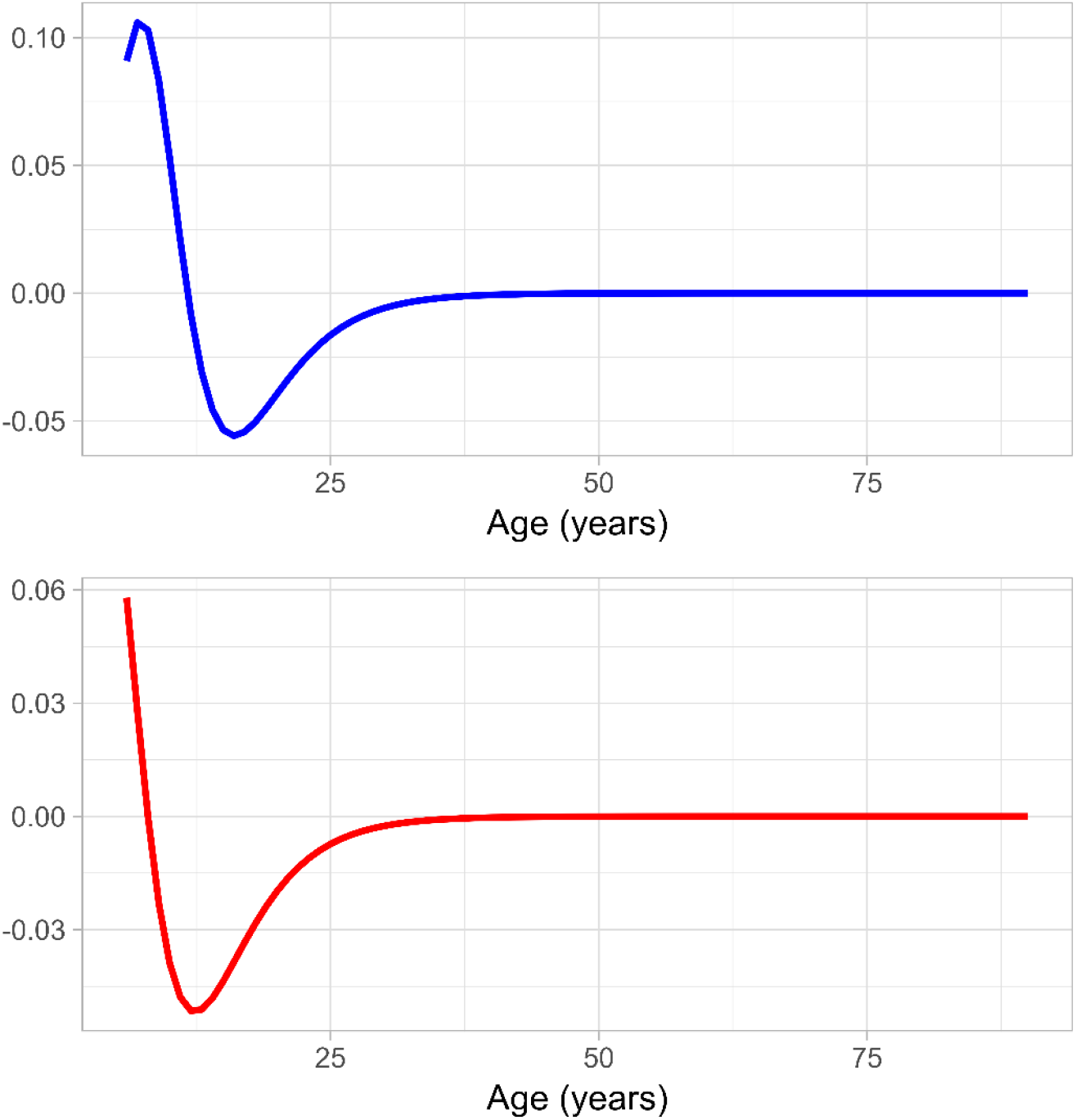
Second Derivative of the Gompertz Model for General and Emotional Vocabulary Growth. The plot displays the second derivative of the fitted Gompertz curve for general (blue dots) and emotional (red dots) vocabulary scores. The second derivative reflects the deceleration of vocabulary acquisition over time. The minimum point indicates the age at which the rate of vocabulary growth is maximized.

Normalized fitted curves for general and emotional vocabulary are displayed in Figure 3. The curves were scaled relative to their respective asymptotes to facilitate direct comparison of growth dynamics. The normalized trajectory for emotional vocabulary showed a more rapid rise and reached near-asymptotic levels earlier in development compared to general vocabulary. In contrast, the general vocabulary curve indicated a slower and delayed growth pattern, with vocabulary acquisition extending over a broader age range.

**Figure 3.**
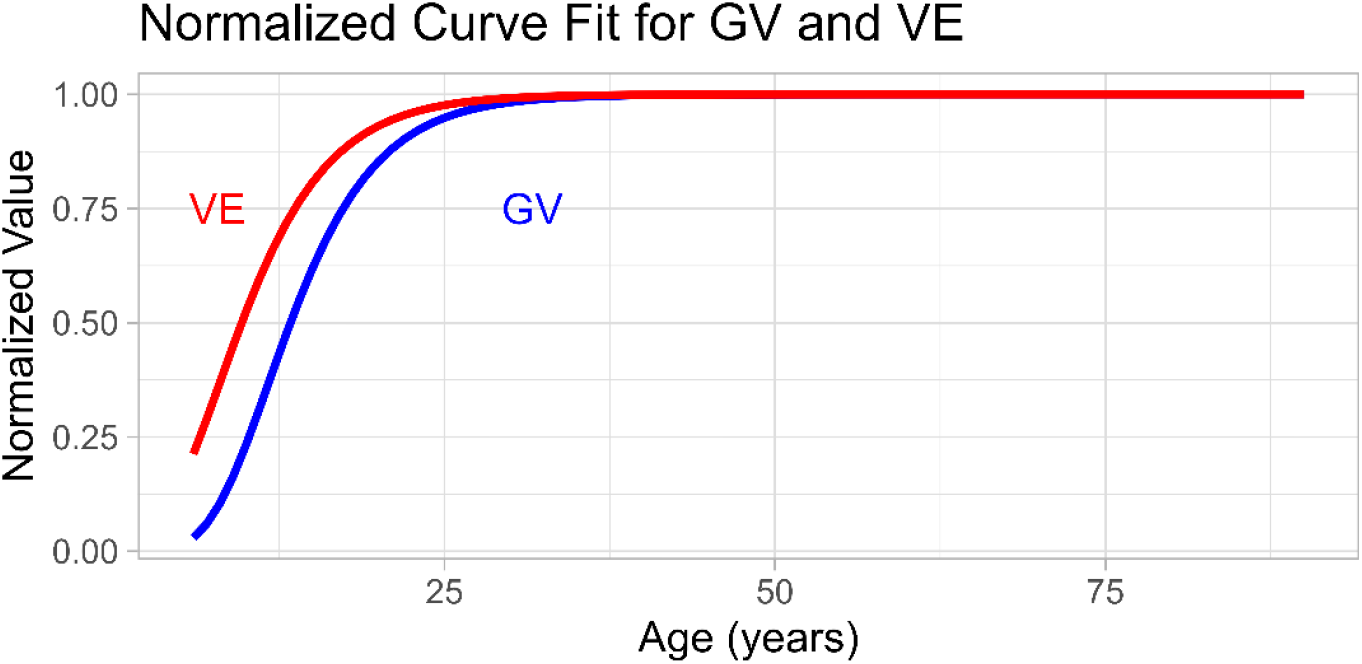
Normalized Gompertz Growth Curves for General and Emotional Vocabulary. The plot shows normalized fitted curves for general vocabulary (GV, blue line) and emotional vocabulary (VE, red line). Values were scaled relative to each model’s asymptote to allow direct comparison. Emotional vocabulary exhibited earlier and faster growth compared to general vocabulary.

To directly compare the growth trajectories of GV and EV, a joint Gompertz model incorporating interaction terms between vocabulary type and growth parameters was fitted (table 3). The results showed a significant positive adjustment in the inflection point (Coef_C = 3.71, p < .001), indicating that GV develops later than EV.

**Table 3.**
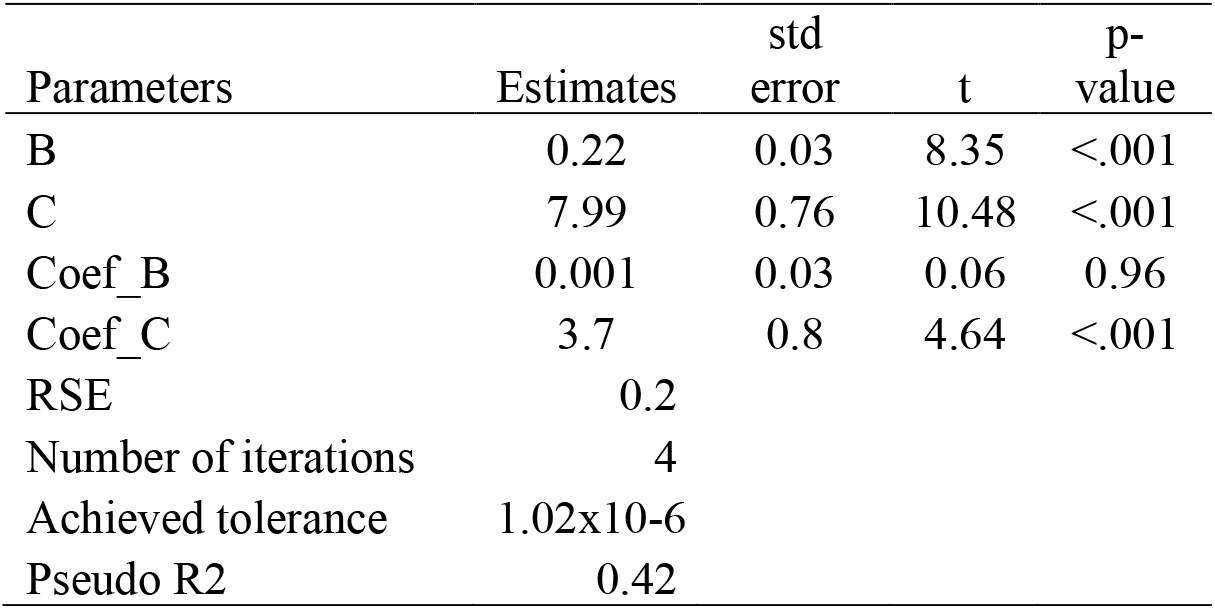
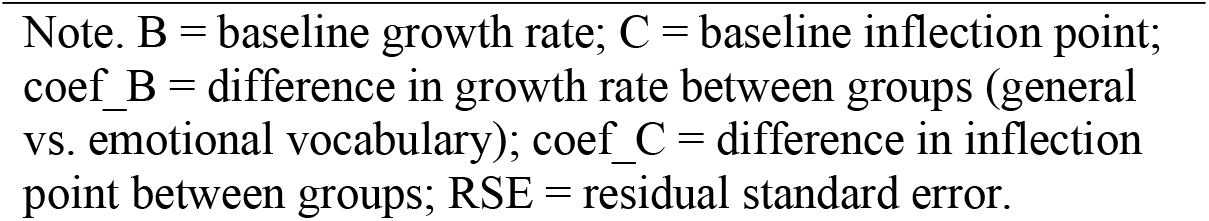
Parameter Estimates and Model Fit Statistics for the Joint Gompertz Model Comparing General and Emotional Vocabulary Growth.

These findings were corroborated by a nonlinear mixed-effects model, which accounted for individual differences by including a random effect on the inflection point parameter. Fixed effects estimates closely matched those obtained in the non-linear least squares model (Coef_C = 3.70, p < .001), and the standard deviation of random effects was small, suggesting limited individual variability.

As a complementary analysis, a linear mixed-effects model was fitted to normalized vocabulary scores with fixed effects of vocabulary type (emotional vs. general), age (each individual age), and their interaction, and random intercepts for individuals. The model revealed significant main effects of vocabulary type (F_(1,784)_ =229.49, p < .001) and age group (F_(35,784)_ = 20.59, p < .001), as well as a significant interaction between vocabulary type and age group (F_(35,784)_ = 3.79, p < .001). Post hoc pairwise comparisons indicated that emotional vocabulary scores were significantly higher than general vocabulary scores during adolescence and early adulthood— between 12 and 20 years of age—but no significant differences were observed among older adults (Figure 5). These results suggest that the advantage of emotional vocabulary development is prominent earlier in life but converges with general vocabulary in later adulthood.

Figure 4 displays mean normalized vocabulary scores and standard errors for emotional and general vocabulary across age. The figure visually confirms the patterns observed in the model-based analyses: emotional vocabulary scores were consistently higher than general vocabulary scores from adolescence through adulthood. In older age groups, the gap between emotional and general vocabulary narrowed, and greater variability was observed, likely reflecting reduced sample size among older participants. Overall, the plot supports the conclusion that emotional vocabulary develops earlier and more rapidly, with general vocabulary reaching similar levels at later ages.

**Figure 4.**
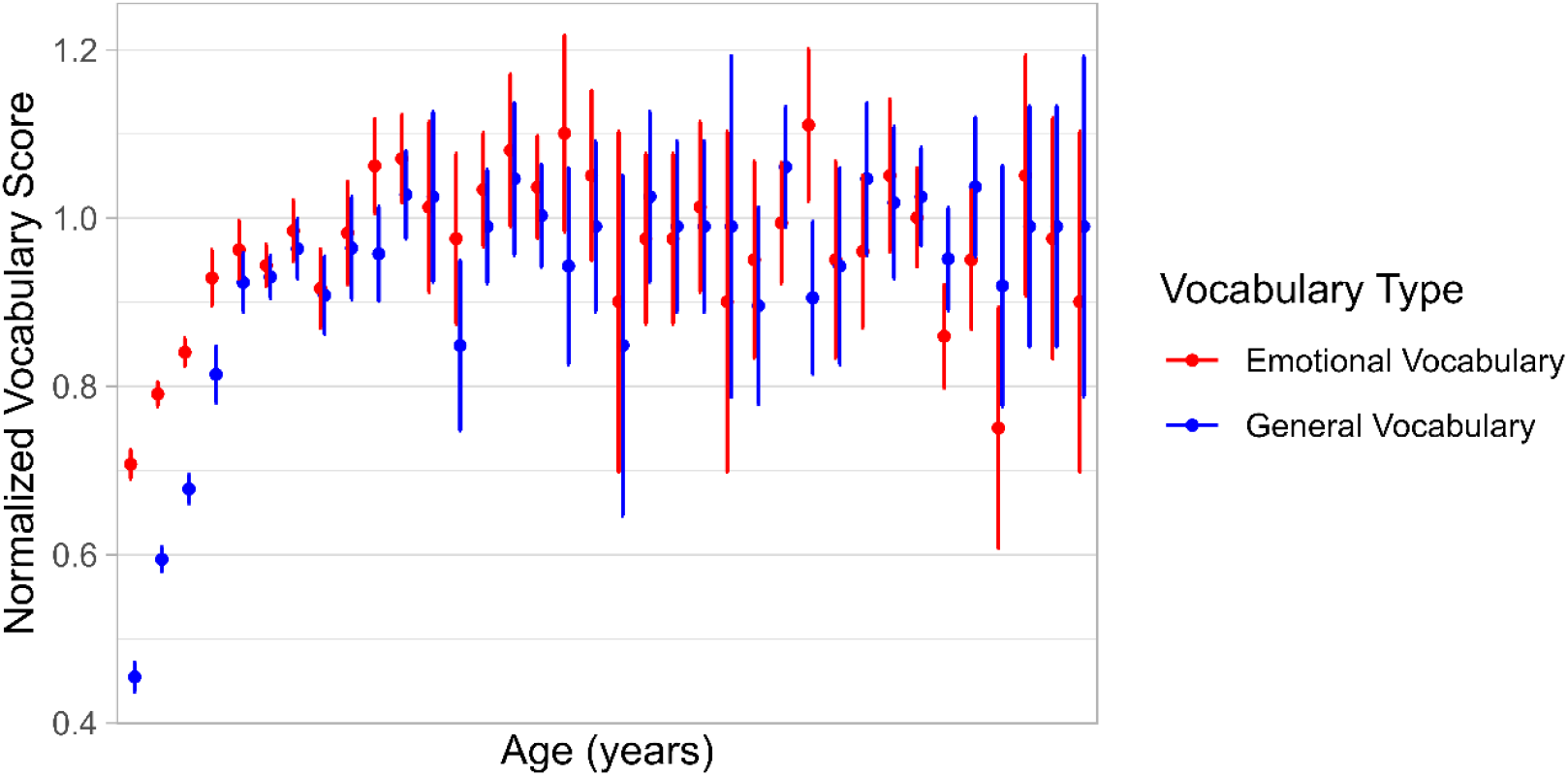
Mean Normalized Vocabulary Scores for Emotional and General Vocabulary by Age. Mean normalized vocabulary scores for emotional (red) and general (blue) vocabulary are plotted across age groups. Error bars represent ±1 standard error of the mean. Emotional vocabulary scores were consistently higher than general vocabulary scores in younger participants, with differences diminishing in older age groups. Greater variability in scores is observed at older ages, reflecting smaller sample sizes.

## Discussion

The present study examined the developmental trajectories of emotional and general vocabulary across the lifespan, modeling growth using Gompertz functions. The results provide compelling evidence for distinct yet convergent patterns of vocabulary acquisition: emotional vocabulary exhibited a faster and earlier growth trajectory, while general vocabulary developed more slowly and reached asymptotic levels later in life.

The use of the Gompertz function to model vocabulary growth proved particularly appropriate. This flexible nonlinear model captured the asymmetric nature of vocabulary development, characterized by early acceleration followed by a progressive plateau (Tjørve & Tjørve, 2017). Importantly, the parameters of the Gompertz model — asymptote, growth rate coefficient, and inflection point — offered interpretable markers aligned with developmental theory: the asymptote reflected the upper limit of vocabulary acquisition, the growth rate captured the pace of learning, and the inflection point indicated the age at which vocabulary growth was maximal. Visual inspections of model fits and second derivative plots provided deeper insight into the developmental process, highlighting how growth accelerates and decelerates with age. The Gompertz function adequately captured the observed trajectories, yielding minimal residual errors and approximately normal distributions of residuals.

The findings regarding emotional vocabulary development are consistent with the Theory of Constructed Emotion (Barrett, 2017a; Barrett, 2017b; Hoemann, Wu, et al., 2020), which posits that emotional concepts are formed through accumulated conceptual knowledge and linguistic experience. In line with this framework, emotional vocabulary in the present study showed rapid acquisition during adolescence and reached near-asymptotic levels in early adulthood. This supports prior evidence suggesting that adolescence is a key period for the refinement and expansion of emotional conceptual knowledge (Bazhydai et al., 2019; Grosse et al., 2021).

Our results further suggest that emotional vocabulary develops earlier than general vocabulary. This pattern may reflect the foundational role that emotional concepts and language play in cognitive and socioemotional development. According to the Theory of Constructed Emotion (Barrett, 2017<u>a</u>), language is essential for organizing sensory inputs into coherent emotional categories, facilitating emotional experience and regulation. Similarly, conceptual development models (Nelson, 2007) emphasize that concepts tied to emotional and social experiences emerge early because of their motivational and adaptive significance. Thus, the earlier acceleration of emotional vocabulary acquisition observed here may reflect an adaptive cognitive priority, enabling individuals to navigate complex emotional and social environments from an early age. The slower and prolonged growth of general vocabulary observed in our study aligns with broader cognitive models that characterize general lexical acquisition as a gradual, experience-dependent process (Cain & Oakhill, 2014; Kintsch, 1998; Ouellette, 2006). This pattern also parallels findings from second-language acquisition research (Song et al., 2015), which suggest that general vocabulary learning continues over extended periods, relying on increasingly sophisticated learning strategies.

The convergence observed between emotional and general vocabulary trajectories in older adulthood provides novel insights. While earlier stages showed a consistent advantage for emotional vocabulary, this gap disappeared in older adults. The increased variability observed among older adults, as evidenced by wider standard errors, may reflect cohort effects, heterogeneous educational experiences, or cognitive aging processes that warrant further investigation.

These findings have theoretical and applied implications. Theoretically, they support the view that emotional vocabulary is an essential cognitive resource, closely tied to emotion regulation, social interaction, and psychological well-being (DeLap et al., 2025; Kashdan et al., 2015; Pond Jr. et al., 2012). Applied, they highlight adolescence as a critical window for interventions aimed at enhancing emotional differentiation and vocabulary sophistication, potentially yielding long-term benefits for emotional health.

Several limitations must be acknowledged. The cross-sectional design restricts the ability to infer individual developmental trajectories definitively. Longitudinal studies are needed to confirm the temporal patterns observed and to assess potential cohort effects, especially in older populations. Additionally, while the sample covered a wide age range, representation of older adults was relatively limited, suggesting the need for larger and more balanced samples in future research.

Despite these limitations, the present study advances understanding of vocabulary development across the lifespan and demonstrates the utility of the Gompertz function for modeling complex, nonlinear growth patterns in cognitive and linguistic domains. By directly comparing emotional and general vocabulary trajectories, the findings contribute to ongoing discussions about the role of language in emotional development and offer a robust empirical basis for future longitudinal and intervention research.

## Data Availability

Data described in the manuscript, code book, and analytic code will be made publicly and freely available without restriction at https://github.com/JFormoso/vocabulary-gompertz and in the permanent repository https://doi.org/10.6084/m9.figshare.29386472.

## Declaration of conflicting interest

The authors declared no potential conflicts of interest with respect to the research, authorship, and/or publication of this article.

## Ethical statement

This study was reviewed by the Responsible Conduct of Research Committee of the Faculty of Psychology at the University of Buenos Aires and the Human Research Ethics committee of the Faculty of Biomedical Sciences, Universidad Austral (P21-072). All participants provided informed consent before taking part in the assessment. Data were anonymized prior to analysis to protect participants’ identities.

During the preparation of this manuscript, the authors used OpenAI’s GPT-3.5 to enhance language clarity and readability. All content generated with this tool was carefully reviewed and edited by the authors, who take full responsibility for the final version of the publication.

## Funding

This work was supported by the National Scientific and Technical Research Council (CONICET) under Grant RESOL-2022-1930-APN-DIR#CONICET (PIBAA), and by the National Agency for the Promotion of Research, Technological Development and Innovation (FONCYT) under Grant RESOL-2023-31-APN-DANPIDTYI#ANPIDTYI (PICT-2021).

## Supplemental material

Supplemental material for this article is available online at https://github.com/JFormoso/vocabulary-gompertz

## Notes

https://github.com/JFormoso/vocabulary-gompertz

### Competing Interest Statement

The authors have declared no competing interest.

